# Expansive pseudogenization and incipient genome degradation of an obligate fruit fly-bacterial mutualism

**DOI:** 10.1101/2025.08.03.667778

**Authors:** Thorsten E. Hansen, Renee L. Corpuz, Tyler J. Simmonds, Charlotte Aldebron, Scott M. Geib, Charles J. Mason, Sheina B. Sim

## Abstract

Obligate microbial symbioses are often characterized by streamlined biosynthetic pathways and reduced genomes. The evolutionary process of this reduction first involves an increase in the abundance of non-functional coding genes (pseudogenes) followed by their removal. The olive fruit fly (*Bactrocera oleae*) harbors an extracellular symbiotic gut bacterium *Candidatus Erwinia dacicola*, which is crucial to its usage of fruit from the olive genus *Olea* as a larval food source. In this study, we combined genomics and transcriptomics of *Ca. E. dacicola* to investigate pathways that facilitate this mutualism. Of 4,675 genes in the *Ca. E. dacicola* genome, 1,783 were classified as pseudogenes. Some biochemical pathways such as amino acid pathways, biofilm regulator BssS, and 6-phospho-β-glucosidase which are implicated in hydrolyzing oleuropein were complete. However, pathways connected to baseline homeostasis, which would impact cellular functions needed for a bacterium to be free-living, were heavily pseudogenized. Gene selection analyses in *Ca. E. dacicola*, when compared to related organisms, indicated positive selection on genes related to amino acid metabolism, carbon utilization, transport, and energy production. Our results indicate that *Ca. E. dacicola* is likely producing amino acids and metabolizing plant phytochemicals. These results reveal that the *Ca. E. dacicola* genome is undergoing incipient genome erosion in support of an unculturable obligate mutualism.

**Importance:** Many beneficial bacteria that live inside insects have highly reduced genomes, but little is known about the transitional stages that occur as free-living microbes evolve into obligate symbionts. We show that the olive fruit fly symbiont, *Candidatus Erwinia dacicola*, retains hallmarks of its plant-associated ancestry while undergoing extensive genome degradation, including the accumulation of mobile DNA elements and inactive genes. At the same time, genes involved in nutrient production, environmental persistence, and host interactions remain functional and are evolving under selection, providing a rare snapshot of how bacterial genomes are reshaped during the evolution of an obligate mutualism. These findings have implications for how obligate gut symbioses are formed and maintained in insect herbivores.

## Introduction

From evolutionary and ecological perspectives, there are genomic features that are frequently associated with obligate microbial symbioses. As a byproduct of different evolutionary forces, these bacteria typically have extremely small genomes, are extremely coding dense, and AT-biased (Burke & Moran, 2011; Moran et al., 2008; Salem et al., 2015; Sudakaran et al., 2017). One process many obligate symbionts are prone to is Muller’s ratchet which predicts that small asexual populations in the absence of recombination are susceptible to accumulating deleterious mutations (Allen et al., 2009; Moran, 2016; Naito & Pawlowska, 2016). In concept, bacteria undergoing a transition from being free-living and stochastic to more intimate host-associated will have genomes with specific characteristics due to accumulated deleterious mutations (McCutcheon & Moran, 2012). These traits include a genome size comparable to free-living relatives, a genome with lost or reduced functions due to an abundance of pseudogenes, and a lower coding density. While there are documented cases of bacteria with these characteristics, most of them are intracellular or have begun to exhibit instances of genome decay. Characterizing these symbioses can provide insight into the evolution of herbivory and co-option of microbial resources by hosts insects.

Most pestiferous fruit flies (Diptera: Tephritidae) are often highly polyphagous herbivores (Dominiak & Follett, 2024; Vargas et al., 2015) and harbor gut microorganisms which contribute to digestion (Hafsi & Delatte, 2023; Raza et al., 2020). The gut microbiome of tephritid fruit flies are often described as facultative commensalisms, which can vary between life stages, populations, and other environmental conditions. Unlike other Tephritidae, the olive fruit fly, *Bactrocera oleae*, is an oligophagous pest of olives (*Olea* spp.) (Daane & Johnson, 2010) and is nearly exclusively colonized by bacterial strains of *Ca. Erwinia dacicola* (Capuzzo et al., 2005).

The olive fruit fly and its symbiosis with *Ca. Erwinia dacicola* has long been investigated. First described in the early 1900s (Petri, 1909), *Ca. E. dacicola* is an extracellular mutualist observed in the gut in larval, pupal, and adult life stages (Siden-Kiamos et al., 2022), and spends a part of its lifecycle exposed when the egg surface is inoculated during oviposition into olive flesh (Capuzzo et al., 2005). Other bacterial taxa can be present in the system, but they are often inconstant and comprise a small fraction of the microbial biomass (Blow et al., 2020; Capuzzo et al., 2005; Koskinioti et al., 2019). *Ca. E. dacicola* has traits that indicate it is an obligate symbiont: it is unculturable, not detected outside its association with the olive fruit fly (Capuzzo et al., 2005), and the host experiences severe negative fitness effects when *Ca. E. dacicola* is experimentally depleted (Ben-Yosef et al., 2010; Ben-Yosef et al., 2015; Ben Yosef et al., 2014; Fytizas & Tzanakakis, 1966; Hagen, 1966; Jose et al., 2019). Olive fly larvae can feed in unripe fruit, and prior research has determined that *Ca. E. dacicola* is crucial for successful host production (Ben-Yosef et al., 2015). Most of the described *Erwinia sp*. are plant pathogens (Kharadi et al., 2021; Marín et al., 2011; Palmer et al., 2017). Besides *Ca. E. dacicola*, *Ca. E. haradaeae* found in aphids*, Ca. E. impunctatus* in midges, and *Erwinia sp*. in Western Flower Thrips are thus far the best characterized insect-associated organisms in the genus (de Vries et al., 2001; de Vries et al., 2004; Manzano-Marı n et al., 2020; Pilgrim, 2024).

Prior work has indicated that *Ca. E. dacicola* has a genome size typical of a free-living microorganism with intact biosynthetic pathways (Blow et al., 2020; Blow et al., 2016; Estes, Hearn, Agrawal, et al., 2018; Estes, Hearn, Nadendla, et al., 2018; Pavlidi et al., 2017). Given that *Ca. E. dacicola* is unculturable, and these prior reports indicate is has an estimated genome size of 2.1 mb – 2.9 mb, 72 – 76% coding density, and 52 – 53% GC content (Blow et al., 2020; Blow et al., 2016; Estes, Hearn, Nadendla, et al., 2018), we hypothesized that the genome represents the evolutionary stage prior to erosion.

We performed meta-transcriptome sequencing and utilized a high-quality and gapless genome of *Ca. E. dacicola* (Hansen et al., 2025) to address three central hypotheses relating to the co-evolution of symbiont and host: 1) *Ca. E. dacicola* shares ancestry with *Erwinia* plant pathogens and will retain plant pathogenic effector genes, 2) *Ca. E. dacicola* is undergoing incipient genome erosion where pseudogenes are caused by the insertion of transposable elements across the genome, and 3) positive selection is acting on genes that are important for host growth and survival in nutritionally limited substrates, such as nutritional provisioning genes linked to amino acid production.

## Results

### Phylogenetic placement

Phylogenomic analysis showed that the UH Lalamilo isolate *Ca. E. dacicola* grouped in a clade with previously published *Ca. E. dacicola* genomes. Sister clades to *Ca. E. dacicola* (i.e., *E. aphidicola*, *E. rhapontici*, *E. sorbitola*, *E. persicina*) are all plant pathogens with high bootstrap values (Supplementary figure 1).

### Effectors in the *Ca. E. dacicola* genome

Bacterial effector proteins can be important for mediating interactions in hosts by bacterial pathogens. As *Ca. E. dacicola* appears to be derived from plant pathogenic bacteria, bacteria effector proteins may be involved between these microbes and the fly host. An effector-specific annotation showed putative effectors spread across the *Ca. E. dacicola* genome (Table S3). In terms of putative effector proteins, 80 total type IV effector proteins were predicted. Effectors across the *Erwinia* plant pathogens (*E. amylovora*, *E. aphidicola,* and *E. persicina*) showed largely distinct orthogroups, with *E. amylovora* having the greatest number, followed by *E. aphidicola,* and *E. persicina* (Figure 1). The amount of effector orthogroups shared between these pathogens was relatively minimal as only the *E. amylovora* - *E. persicina* comparison had more than ten orthogroups (Figure 1). Surprisingly, *Ca. E. dacicola* possessed a small number of effectors orthogroups (i.e., 12) and shared none with the other *Erwinia* plant pathogens (Figure 1).

**Figure 1:**
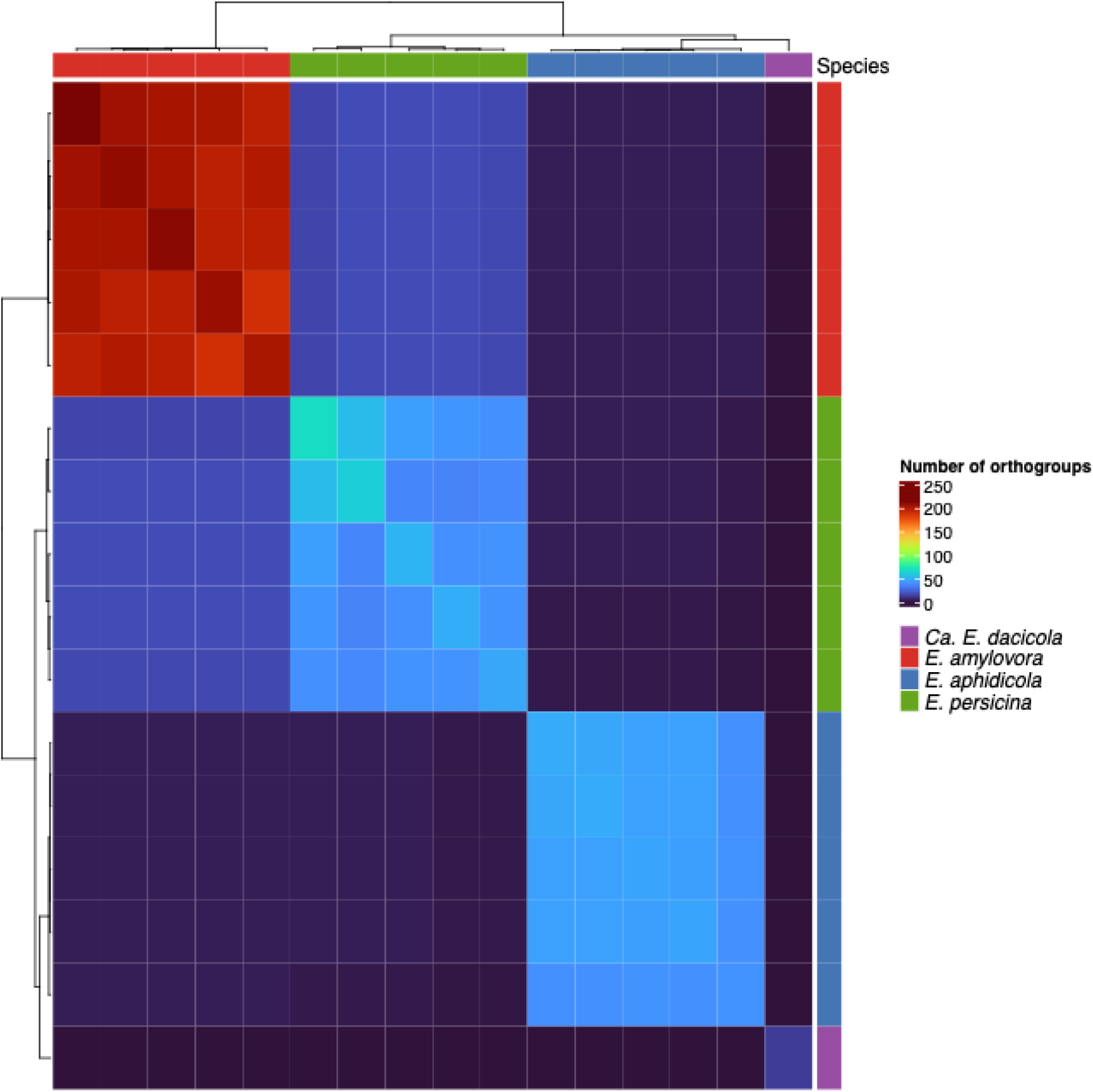
Heatmap representation of distinct and shared effector orthogroups between *Candidatus Erwinia dacicola* UH Lalamilo and *Erwinia amylovora, Erwinia aphidicola, and Erwinia persicina* (i.e., 5 genomes from each of these plant pathogens).

### Incipient genome erosion of *Ca. E. dacicola*

*Ca. E. dacicola’s* genome had a very high number of pseudogenes (1783), comprising 38.13% of the genome. PGAP indicated these pseudogenes were categorized as frameshifts (762), coding for partial proteins (i.e., non-coding) (1279), internal stops (597), multiple problems (697), and a truncated protein (1). Many of these pseudogenes, such as the frameshift and internal stops, were likely caused by transposable elements (TEs). The genome *Ca. E. dacicola* was found to possess a high number of TEs (714) which comprise 15.27% of the genome. The majority of TEs have undergone pseudogenization (494) with 220 still being active coding sequences (Table S4). Pseudogenes were significantly enriched for transposon associations relative to intact genes (Fisher’s exact test, odds ratio = 2.17, 95% CI = 1.59–2.97, p = 4.28 × 10), suggesting that transposon activity may contribute to pseudogene formation in this genome. Most DNA recombination and repair genes were present and complete. However, one DNA recombination and repair gene was entirely absent (Table S5). The loss of DNA repair genes suggests that deleterious mutations are more likely to accumulate over time.

To understand which pathway functions have been impacted by pseudogenes, KEGG KO numbers were assigned to genes, when available, and ORA KEGG analysis was conducted comparing pseudogenes to coding sequences genes. ORA analysis identified several pathways significantly affected by pseudogenization (Table S6). Using transcriptomic data, we observed residual expression of some of these pseudogenes, but usually not greater than expression of coding genes in the same pathways (Figure 2).

**Figure 2:**
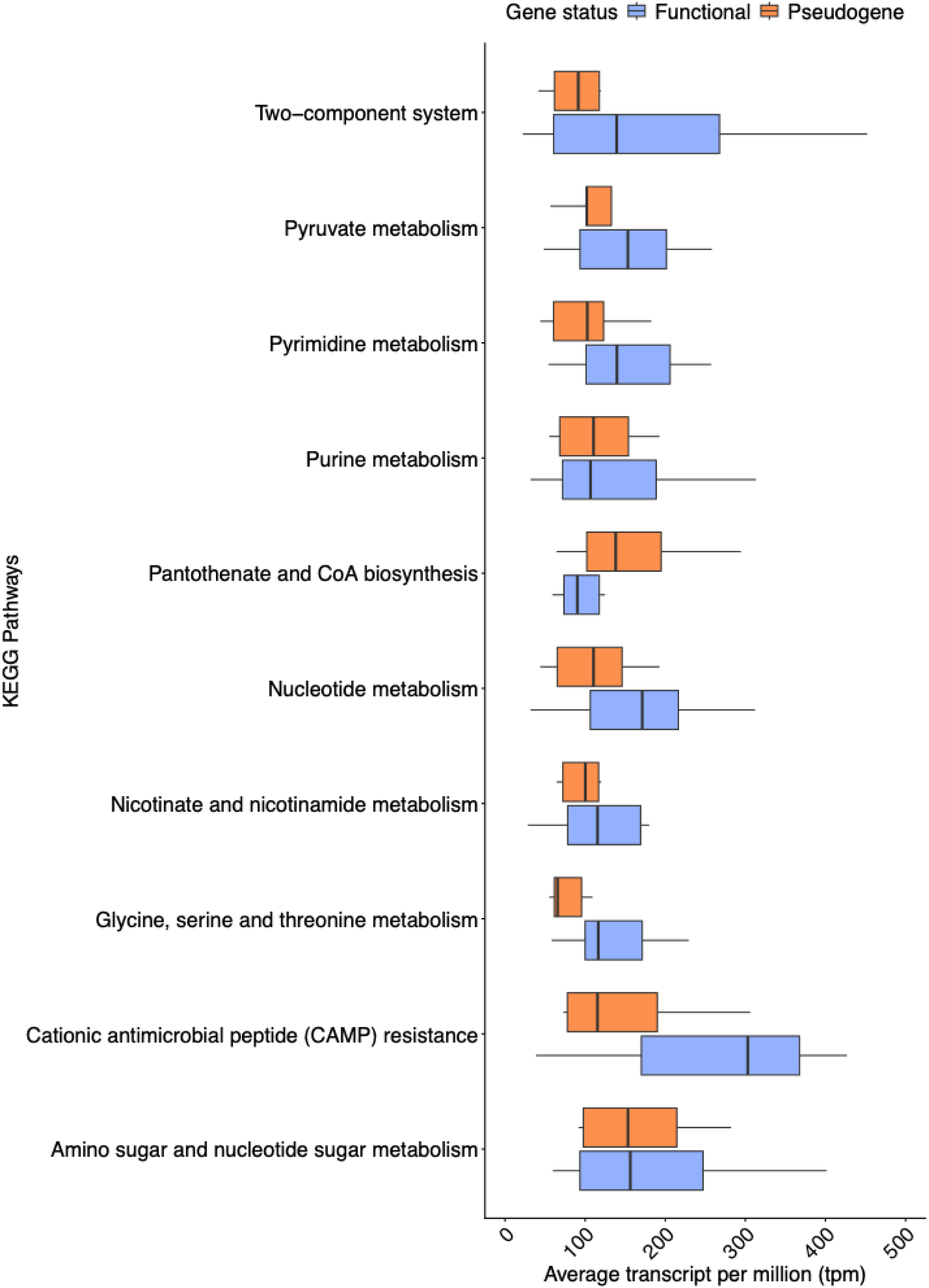
Boxplot showing average transcript per million for gene spread across different KEGG pathways. These data are grouped between genes that are actual functional and those that are pseudogenes. All KEGG pathways shown here are pathways that were statistically signicantly enriched in over representation analysis when comparing functional genes and pseudogenes.

### Pathway completeness

KEGG Decoder was used to assess the minimum number of genes needed for bacterial pathway completeness. Overall, there was general trend of genes connected to biosynthesis (amino acids and cofactors) and central carbon & energy metabolism having the most complete pathways and being least impacted by pseudogenes (Figure 3). Most amino acid pathways appeared complete (TableS7). Components of the pathways to glycine, serine, and threonine may not be fully functional considering that some genes pertaining to these pathways were pseudogenes. Additionally, tryptophan likely can no longer be produced as its pathway is incomplete and has pseudogenes. From a metabolic standpoint, there were complete pathways for sulfur assimilation, enzymes to breakdown starch molecules, and pathways to produce isoprenoids (Table S7). There were also complete pathways for oxidative phosphorylation and ATP production (Table S7). *Ca. E. dacicola* also has the capacity to produce riboflavin *de novo* from GTP and ribulose-5P (Table S7). It encodes complete pathways for many transporters: ammonia, copper, Fe-Mn, ferrous iron, phosphate, and thiamin (Table S7). The Sec-SRP pathway and Biofilm regulator BssS pathway both hint at larger environmental interactions.

**Figure 3:**
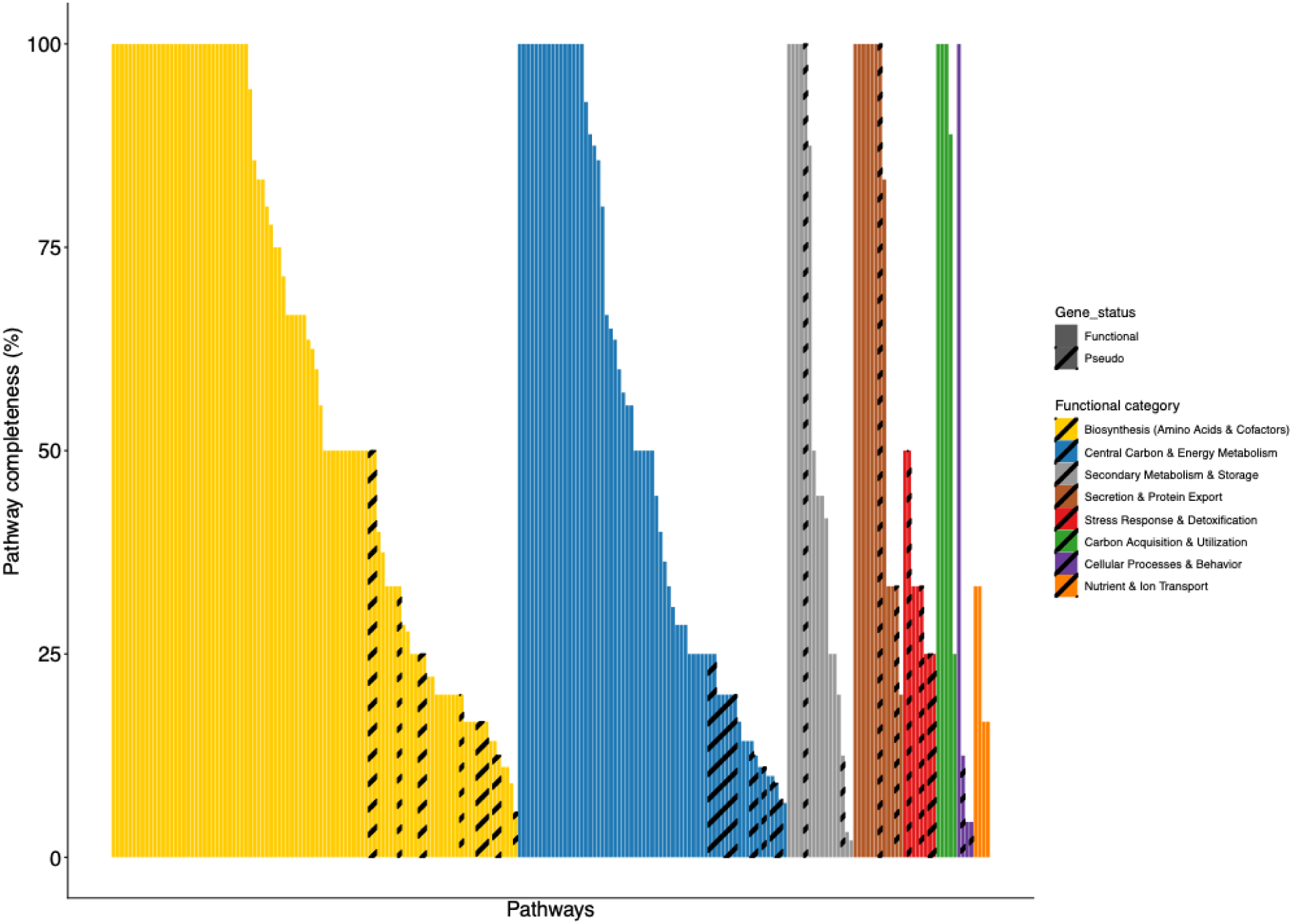
Barplot representation of percent completion of pathways, in the *Candidatus Erwinia dacicola* genome, across functional categories and ranked within categories by decreasing completeness. The pathways shown are KEGG pathways and additional, non-KEGG, pathways which were assed using a small targeted HMM database.

Many pathways connected to glycolysis and energy production were incomplete (Table S7), while carbon fixation and fermentation may have reduced function or be entirely lost. Secretion systems were another large category of incomplete pathways. There are also incomplete pathways for some detoxification, specifically arsenic reduction and formaldehyde assimilation. *Ca. E. dacicola* was also found to have incomplete chemotaxis and flagellum pathways. The last incomplete pathway found was NADH-quinone oxidoreductase (oxidative phosphorylation).

### Positive selection on genes linked to symbiosis

To identify signatures of positive selection on *Ca. E. dacicola* genes, dN/dS ratios of single copy orthologs were compared between other *Erwinia* and *Ca. E. dacicola* with distinct lifestyles, *E. amylovora* and *E. persicina* (extracellular, free-living plant pathogen), *E. aphidicola* (plant pathogen and insect associate) and *Ca. E. haradaeae* (intracellular aphid symbiont). The total number of single copy orthogroups for each comparison were: 946 in *E. amylovora*, 945 in *E. persicina*, 958 in *E. aphidicola*, and 95 in *Ca. E. haradaeae*. Each branch model analysis only tested a one-to-one species comparison. To see if selection was acting similarly across these different *Ca. E. dacicola* to pathogen comparisons each analysis was filtered to only contain the single copy orthologues held in common. Additionally, any dN/dS values for the foreground branch (i.e., *Ca. E. dacicola*) that were below 0.1 or above 3.0 were removed. This range was selected as value below or above it reflects sequences too similar or two divergent to compare. After both trimming steps, there were 112 genes across all three *E. species* pathogen comparisons (Table S8) and 17 total genes for the *Ca. E. haradaeae* comparison (Table S9) undergoing positive selection.

In comparison to *E. amylovora*, *E. aphidicola,* and *E. persicina*, there were many pathways which had genes under positive selection (Figure 4). Several genes under positive selection were connected to nutritional pathways, such as amino acid production (acetylornithine deacetylase. ω = 1.38356; branched-chain amino acid transaminase, ω = 1.98933; diaminopimelate epimerase, ω = 1.05223; NADP-dependent phosphogluconate dehydrogenase, ω = 1.02745; 4-hydroxy-tetrahydrodipicolinate reductase, ω = 1.20633) and vitamin synthesis (branched-chain amino acid transaminase, ω = 1.98933; bifunctional UDP-sugar hydrolase/5’-nucleotidase UshA, ω = 1.21503; pyridoxamine 5’-phosphate oxidase, ω = 1.21637). Environmental interaction and transporters were also prominent and under positive selection (branched-chain amino acid transaminase, ω = 1.98933; phosphate ABC transporter ATP-binding protein PstB, ω = 1.04627; phospholipid ABC transporter ATP-binding protein MlaF, ω = 1.59554; two-component system sensor histidine kinase EnvZ, ω = 2.51219; 4-hydroxy-tetrahydrodipicolinate reductase, ω = 1.20633)

**Figure 4:**
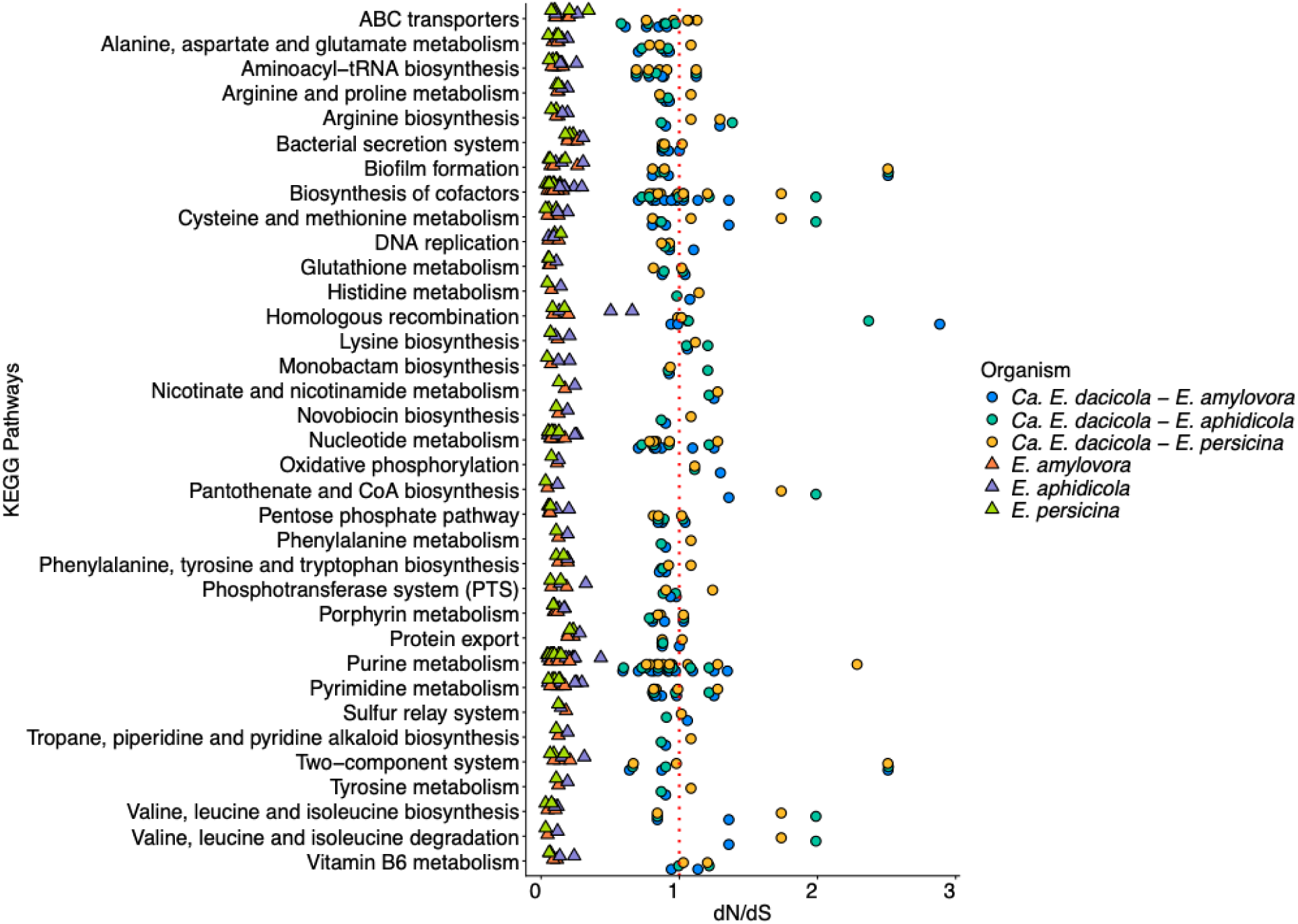
Boxplot for the dN/dS values of shared single copy ortholog genes between *Candidatus Erwinia dacicola* and *Erwinia amylovora, Erwinia aphidicola, and Erwinia persicina*. The dN/dS values showed between *Candidatus Erwinia dacicola* and the three separate Erwinia sp. correspond to individual analyses between each foreground and background species comparison. Genes were grouped by KEGG pathways. Some genes have multiple ko terms and are associated with multiple pathways. A red dashed line is set at dN/dS = 1 to better visualize all genes experiencing positive selection.

In comparison to *Ca. E. haradaeae*, the largest set of genes under positive selection were related to the electron transport chain and oxidative phosphorylation (electron transport complex subunit RsxA, ω = 1.30723; electron transport complex subunit RsxG, ω = 1.00605; NADH-quinone oxidoreductase subunit NuoG, ω = 1.20255) (Figure 5). Aminoacyl-tRNA biosynthesis (phenylalanine--tRNA ligase subunit alpha, ω = 1.1728) and ribosome biogenesis (23S rRNA pseudouridine (955/2504/2580) synthase RluC, ω = 1.20496) were under positive selection (Figure 4). A few genes were likely under stabilizing selection: DNA replication (replicative DNA helicase, ω = 1.00939), glutathione metabolism and pentose phosphate pathway (NADP-dependent phosphogluconate dehydrogenase, ω = 1.01726), and purine metabolism (glutamine-hydrolyzing GMP synthase, ω = 1.01631).

**Figure 5:**
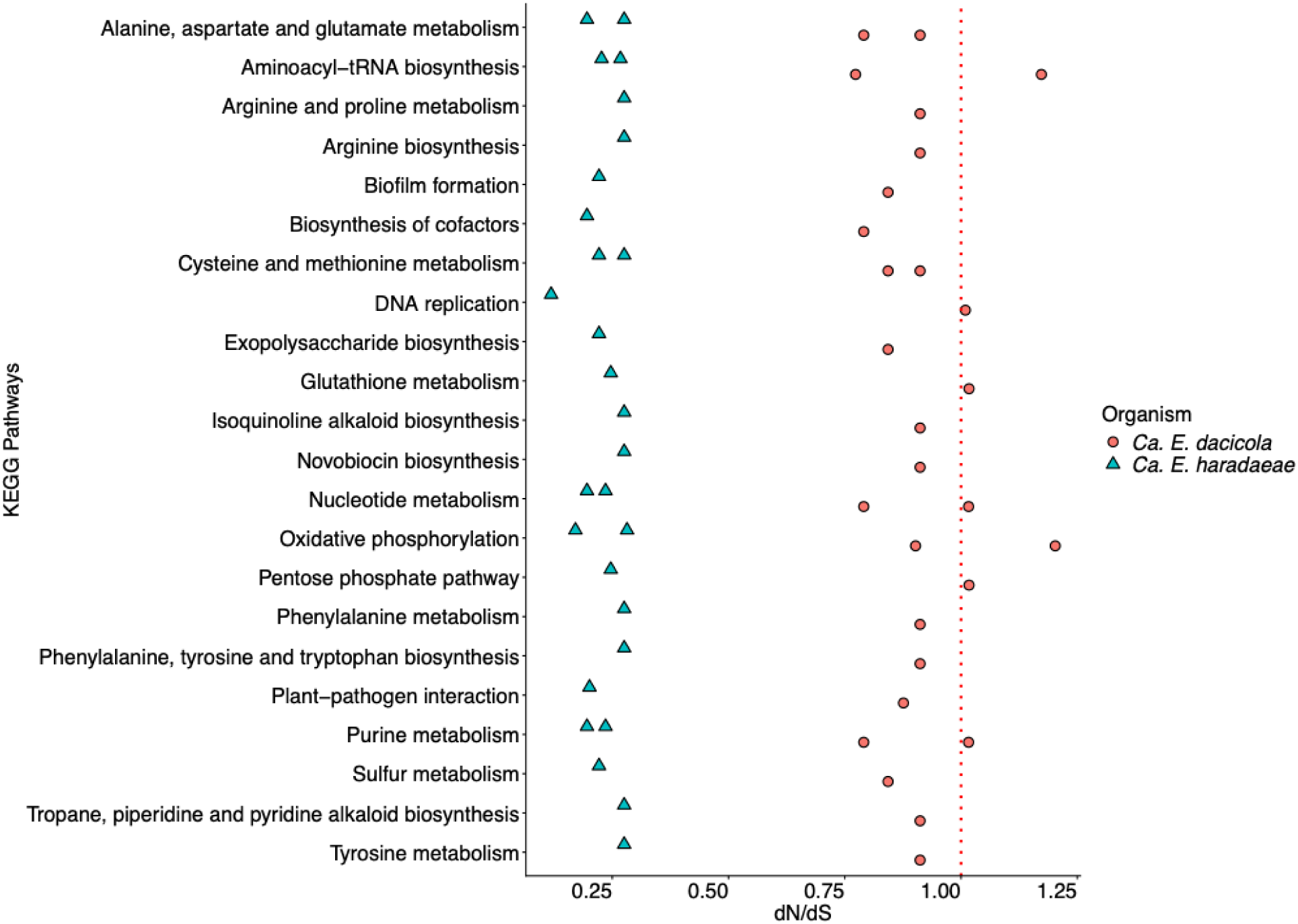
Boxplot for the dN/dS values of shared single copy ortholog genes between *Candidatus Erwinia dacicola* and *Candidatus Erwinia haradaeae*. Genes were grouped by KEGG pathways. Some genes have multiple ko terms and are associated with multiple pathways. A red dashed line is set at dN/dS = 1 to better visualize all genes experiencing positive selection.

## Discussion

### Genomic architecture of *Ca. Erwinia dacicola* as it relates to other insect symbionts

Like all animals, insects have integrated relationships with microbial symbionts (McFall-Ngai et al., 2013). From a genomic perspective, obligate bacterial symbionts are often reduced in size, are AT-biased, exhibit high coding density and codon reassignment, gene loss, and rapid sequence evolution (McCutcheon & Moran, 2012). These likely arise from strong selection pressures exerted by the host environment such as host immunity and gut physiology; as well as Mueller’s ratchet where deleterious mutations accumulate irreversibly in asexual populations because there is little or no recombination to recreate mutation-free genomes (Pettersson & Berg, 2007) Most examples of obligate relationships are at the point where the host and the symbiont are interconnected and mutually dependent, but there are fewer examples of an intermediate relationship or transitional form (Moran et al., 1993). An intermediate stage would feature little bacterial genome contraction, minimal AT-bias, low coding density, and a higher density of pseudogenes and mobile elements (Burke & Moran, 2011; Garber et al., 2021; Koga & Moran, 2014; Lamelas et al., 2011).

*Ca. E. dacicola* is an obligate gut symbiont in the incipient stages of genome erosion. An important indicator is the enormous number of pseudogenes present in the *Ca. E. dacicola* genome (35% of coding density). Other systems which have “transitional” symbionts have been reported with fewer pseudogene densities (10-25%) (Santos-Garcia et al., 2017). *Ca. Symbiopectobacterium endolongipinus* and *Ca. Sodalis endolongispinus* are similar to *Ca. E. Dacicola* in their proportion of pseudogenized genes (∼45%) (Garber et al., 2021), but are extracellular rather than intracellular (Pantidi et al., 2026; Siden-Kiamos et al., 2022). Transposable elements (TEs) have been suggested to be important in streamlining large genomes for obligate symbionts (Siguier et al., 2014; Siguier et al., 2015). If a bacterium that has adopted a host-restricted lifestyle acquires TEs, there can be rapid proliferations of TEs throughout their genome due to population isolation and a decrease in the strength of purifying selection. The *Ca. E. dacicola* genome possesses a notably large number of TEs (714; 15.27% of coding density) compared to other symbionts such as *Candidatus Sodalis baculum* and *Serratia symbiotica* SCc where none present (Lamelas et al., 2011), (Santos-Garcia et al., 2017) and *Symbiopectobacterium endolongispinus* and *Sodalis Endolongispinus* which have many fewer TEs (50 and 220 respectively) (Garber et al., 2021). Despite these indicators of erosion, *Ca. E. dacicola* has functional DNA recombination and repair genes which likely allow the symbiont to retain its relatively large size.

Although sequencing and mis-assembly errors could exist and potential generate false pseudogene signal (Cooley & Wright, 2024), we see no evidence of this given the high consensus accuracy and low number of pseudogenes in the corresponding olive fruit fly genome (5.8%) relative to those in *Ca. E. dacicola* does not imply sequence errors (Hansen et al., 2025). Furthermore, HiFi reads offer increased utility over short-read methods since they can reveal frameshifts and other genomic rearrangement caused by TEs. Most of the previously published *Ca. E. dacicola* genomes were sequenced with short-read methods and, thus, are much smaller and discontinuous (Blow et al., 2016; Estes, Hearn, Nadendla, et al., 2018). Blow et al. (2020) reported on TEs but were unable to reassemble repetitive mobile elements with the use of short-read sequencing.

### *Ca. E. dacicola* evolution in relation to other insect symbionts and pathogens

Members of *Erwinia* are most often associated with plants (Palmer et al., 2017), many of which are notorious plant pathogens (Kharadi et al., 2021), but there are insect-associated members of this clade. *Ca. E. haradaeae* was recently discovered to have replaced an obligate intracellular symbiont in a complex of different *Cinara* aphids, (Manzano-Marın et al., 2020). *Erwinia* has also been shown to associate with the Highland midge and Western Flower Thrips (de Vries et al., 2001; de Vries et al., 2004; Pilgrim, 2024). Phylogenetically, *Ca. E. dacicola* fits within a clade amongst plant pathogens most closely associated with *E. aphidicola*. This suggests that *Ca. E. dacicola* is ancestrally related to a plant pathogen only recently acquired from the environment and transitioned to a symbiotic lifestyle. There is evidence of this in other systems, as bacterial gut associates of plant-feeding insects include taxa that can be intimately associated with plants (Kaltenpoth & Flórez, 2020; Pirttilä et al., 2023), including other tephritid fruit flies (Cárdenas Hernández et al., 2023; Hendrycks et al., 2022; Morrow et al., 2015; Tian et al., 2023).

The presence of bacterial effector proteins in the *Ca. E. dacicola* genome also hints at its previously pathogenic lifestyle. Bacterial effector proteins modify the host to create conditions favoring bacterial survival (Ellis & Machner, 2024). However, contrary to our first hypothesis, *Ca. E. dacicola* did not share orthologous effector genes with other plant pathogenetic *Erwinia sp*. Instead, the predicted effectors were unique to it in comparison to plant pathogens. This suggests that the *Ca. E. dacicola* effectors are associated with maintaining and regulating associations with its arthropod host rather than mediating plant-based responses at the fruit-egg interface. It is possible these putative effectors in *Ca. E. dacicola* could manipulate host signal transduction or suppress host immune and defense responses (Frank, 2019; Martyn et al., 2022; Miwa & Okazaki, 2017). However, further manipulative experiments should be conducted to verify their functions and impact. Alternatively, since *Ca. E. dacicola* spends a portion of its lifecycle outside of the gut on the egg chorion and in direct contact with plant cells, these genes may be involved in survival during this two-to-three-day period before larval hatch and colonization of host tissues.

### Lost and functional pathways in relation to symbiosis

Previous studies provide evidence that *Ca. E. dacicola* is provisioning nutrients to the host (Blow et al., 2020; Estes, Hearn, Agrawal, et al., 2018), despite the fact that specific pathways like glycine, serine, and threonine metabolism may be nonfunctional due to pseudogenesis. Our data show *Ca. E. dacicola* retaining mostly complete essential and non-essential amino acid pathways which likely help with being extracellular, occupying different organs with the host, and surviving on the surface of the egg during vertical transmission (Estes et al., 2009; Siden-Kiamos et al., 2022). Positive selection on genes involved in amino acid and vitamin synthesis support putative nutritional roles of *Ca. E. dacicola*. In addition, stabilizing selection on glutamine-hydrolyzing GMP synthase and NADP-dependent phosphogluconate dehydrogenase implies a greater role for interconversion of amino acids, ammonia recycling, and the maintenance of glutamine and glutamate (GOGAT cycle) (Hansen & Moran, 2011; Hansen et al., 2020; Price et al., 2014). There was also positive selection on phosphate ABC transporters, which have been implicated in transport of amino acids (Lewis et al., 2012; Tanaka et al., 2018). Positive selection on NADH-quinone oxidoreductase subunit NuoG implies high ATP requirements, which is probably driven by nutrient production (i.e., amino acids) and transport (Cooper, 2000). Furthermore, positive selection on aspartate – tRNA ligase and phenylalanine – tRNA ligase subunit alpha, suggests increased ATP production is needed for increased amino acid synthesis.

Previous research has shown that *Ca. E. dacicola* resides in the guts of adult flies as a biofilm (Estes et al., 2009; Siden-Kiamos et al., 2022). Formation of biofilms may be integral for symbiont survival through pupation and in harsher environments such as the egg surface in oviposition. Our research supports this as the biofilm regulator BssS is complete. Furthermore, we observed positive selection was acting upon the two-component system sensor histidine kinase EnvZ, which responds to changes in osmolarity and is connected to the transition from the reversible to the irreversible attachment phase in biofilm development (Prüß, 2017). In addition to requiring intact genetic pathways that facilitate free-living, *Ca. E. dacicola* also has intact genetic pathways that facilitate the fly host metabolism of harmful phytochemical defense compounds like oleuropein

Olives are laden with phytochemicals, including phenolics, that can be at exceptionally high concentration in unripe tissues. Oleuropein, a glycosylated seco-iridoid phenolic is a prevalent compound that is an antinutrient (Felton & Gatehouse, 1996; Konno et al., 1999; Marra et al., 2020; Spadafora et al., 2008) binding to and inactivating enzymes and reducing protein accessibility (Felton & Gatehouse, 1996; Kroll et al., 2003; Pentzold et al., 2014). It has been proposed, but not metabolically established, that *Ca. E. dacicola* assists in overcoming the oleuropein in olives due to increased larval fitness when feeding on unripe olives (Ben-Yosef et al., 2015). Our assembly indicates that *Ca. E. dacicola* encodes the gene for 6-phospho-β-glucosidase, which has been directly implicated in hydrolyzing oleuropein as seen in *Lactiplantibacillus plantarum* strains (Vaccalluzzo et al., 2022). Esterases have also been implicated in the oleuropein degradation (Ramírez et al., 2017), specifically in tandem with 6-phospho-β-glucosidase (Charoenprasert & Mitchell, 2012; De Leonardis et al., 2016; Vaccalluzzo et al., 2022). Two genes encoding esterases found in the *Ca. E. dacicola* genome, suggesting the potential to metabolize oleuropein to benefit the host.

Although *Ca. E. dacicola* spends a portion of its lifecycle on the egg surface outside of the digestive system, ORA analysis indicated several pathways were impacted thus indicating host restriction. ORA analysis discriminating functional genes and pseudogenes results indicated that baseline homeostasis is heavily impacted, directly affecting many cellular functions needed for a bacterium to be free-living (Fujita et al., 2007; Keweloh & Heipieper, 1996). Furthermore, many physiological processes that are associated with host gut function such as fermentation and sugar utilization have been lost (Goncheva et al., 2022; Paulini et al., 2022). Curiously, even though are *Ca. E. dacicola* extracellular, the regulation and response to environmental conditions have been impaired, including sensing of environmental stimuli and signal transduction, which are important traits for extracellular organisms (Alvarez & Georgellis, 2022; Brown et al., 2022; Golby et al., 1999; Meng et al., 2021; Mensa et al., 2021; Pasqua et al., 2019; Wösten et al., 2017).

## Conclusions

The numerous transposons dispersed across the genome, alongside the high density of pseudogenes, indicate that *Ca. E. dacicola* is in its evolutionary trajectory that follows obligate symbiosis but before widespread genome decay (Bennett & Moran, 2015). Of the complete and fully functional pathways, a majority are associated with host nutrient and allelochemical metabolism and supporting the host-microbe symbiosis. Altogether, there was likely an ancestorial clade of *Ca. E. dacicola* that was plant associated and eventually co-opted by the olive fruit fly but has since lost any plant pathogen effector genes. This is in congruence with estimated evolutionary histories as the olive fruit fly separated from the other species in *Bactrocera* approximately 25 million years ago (Zhang et al., 2022) and has been specializing on olive for ∼ 4-6 million years (Nardi et al., 2005; Teixeira da Costa et al., 2024). Notably, the olive fruit fly – *Ca. E. dacicola* symbiosis differs from other flies in the *Bactrocera* genus, which have more facultative relationships with gut microbiota (Aoki et al., 2025; Cheng et al., 2017; Kempraj et al., 2024; Majumder et al., 2020). Genetic evidence supports systematic changes in the genome have led to a more sessile and commensal bacterium, which is conductive to the mutualistic relationship we currently observe.

## Materials and Methods

### Source material

Infested fruits of *Oleae europaea* were collected at the University of Hawai’i Lalamilo Research Station in Hawai’i County, Hawai’i, USA (<u>20.019814 N, 155.677155 W</u>) and brought to the Daniel K. Inouye United States Pacific Basin Agricultural Research Center in Hilo, Hawai’i, USA (United States Department of Agriculture, Agricultural Research Service). Olive fruits were held for larval development and pupation, and adults were maintained in the laboratory and provided water and food ad libitum (3:1 mixture of granulated sugar and yeast hydrolysate). Eight adult female flies were also taken from this collection and flash frozen in liquid nitrogen for transcriptomic library preparation.

### Transcriptomic library preparation and sequencing

Total RNA was isolated using Direct-zol-96 MagBead RNA kit (Zymo Research) from eight individual female olive fruit flies. Prior to library construction, several quality control steps were performed including quantifying extracted RNA using Qubit RNA BR Assay Kit and the fluorometer function on a DeNovix DS-11 FX+ and assessing RNA quality using the spectrophotometer function on a DeNovix DS-11 FX+ to detect the presence of nucleic acid impurities and the Agilent TapeStation RNA ScreenTape Analysis to determine if there is evidence of sample degradation. Total RNA-seq libraries were constructed using Zymo-Seq RiboFree Total RNA Library Kit (Zymo Research) and sequenced on an Element Biosciences AVITI using a 2×150 Cloudbreak Freestyle High Output sequencing kit. The resulting transcriptome dataset was comprised of ∼ 1316.25 Gb of total host data and ∼ 28.26 Gb of total symbiont data. The percent recovery specifically for symbiont reads ranged from 0.4029 % - 4.0370 % (Table S1).

### *Ca. E. dacicola* genome annotation

Annotation data from the previously published *Ca. E. dacicola* genome (NCBI Bioproject ID <u>PRJNA1090968</u>) was used in this study (Hansen et al., 2025). These annotations come from the NCBI Prokaryotic Genome Annotation Pipeline (PGAP) which predicts protein-coding genes, structural RNAs, tRNAs, small RNAs, and pseudogenes (Tatusova et al., 2016). Additional annotation was done using the GhostKOALA (v 3.1)(Kanehisa et al., 2016) automatic annotation server, which specializes in genome and metagenomic sequences. Here, KO (KEGG Orthology) assignments are performed to characterize gene functions and reconstruct KEGG pathways, BRITE hierarchies, and KEGG modules.

Further, specific annotation was conducted for effectors and transposons. For effector predication, we used the program DeepSecE (v 0.1.1)(Zhang et al., 2023) which uses a deep-learning-based framework for the simultaneous inference of multiple distinct groups of secreted proteins produced by Gram-negative bacteria. DeepSecE can predict type I-IV and VI secretion systems. Transposons were annotated using TnCentral (Ross et al., 2021). TnCentral is a database for prokaryotic transposable elements (TE) which contains approximately 400 annotated TE, including TE from various families, compound transposon, integrons, and associated insertion sequences.

### Phylogenomic analysis

Phylogenomic analysis was conducted on the clade of *Erwinia* (Table S2) using all publicly available *Erwinia* GenBank genomes in the National Center for Biotechnology Information (NCBI). Outgroup genomes were selected based on a previous study on the phylogenomic analysis of the *Pantoea*, *Erwinia*, and *Tatumella* genera (Palmer et al., 2017). Phylogenomic analysis was performed using the pipeline VBCG (v 1.0) (Tian & Imanian, 2023). The VBCG pipeline compares input genomes to the VBGC core gene set, which are twenty core bacterial genes validated across 11,262 species and covering 30,522 complete genomes. A final alignment file, in PHYLIP format, was produced by the pipeline for use. This PHYLIP file was subjected to maximum-likelihood (ML) phylogenetic analysis.

The ML phylogenetic analysis was conducted using RAxML-NG (v 2.0.1) (Kozlov et al., 2019). Analyses were conducted under the JTT amino-acid substitution model with gamma-distributed rate heterogeneity among sites (JTT+G). Tree search and nonparametric bootstrap analyses were performed in a single run using the `--all` workflow. Bootstrap support was assessed using 1000 bootstrap replicates, and both Felsenstein bootstrap proportions (FBP) and transfer bootstrap expectation (TBE) support values were calculated. Tree files produced by RAxML were further annotated using the tree drawing webtool Interactive Tree of Life (iTOL) (Letunic & Bork, 2024).

### Orthologues effectors analysis

To determine if *Ca. E. dacicola* retained pathogenic plant effectors along the course of its evolution, we performed an analysis for orthology. We used DeepSecE (v 0.1.1) to annotate effectors across *Ca. E. dacicola*, *E. amylovora*, *E. aphidicola*, and *E. persicina* (Table S2). The predicted effector amino acid sequences were then used to identify gene orthogroups through OrthoFinder (v 2.5.5)(Emms & Kelly, 2019).

### *Ca. E. dacicola* pathway completeness

The completeness of various metabolic pathways was determined with the kegg-pathways-completeness-tool (v 1.4.0), available through the EMBL European Bioinformatics Institute MGnify platform (Richardson et al., 2023). Using the KEGG KO annotations from GhostKOALA, kegg-pathways-completeness-tool shows percent completeness of established KEGG pathways.

### *Ca. E. dacicola* transcriptomic verification of pseudogenes

Due to a high number of pseudogenes identified with Prokaryotic Gene Annotation Pipeline (PGAP) (see Results), we evaluated the transcription of these genes in the *Ca. E. dacicola* genome. Sample demultiplexing was performed using bases2fastq (v2.0.0.1379264253) (https://docs.elembio.io/docs/bases2fastq/). Initial read quality assessment was performed, for each individual paired read using FastQC (v 0.12.1)(Andrews, 2010) and total quality report was composed with MultiQC (v 1.15)(Ewels et al., 2016). Next, we aligned the RNA-seq reads to the *Ca. E. dacicola* genome to isolate all bacterial reads with BBMap implemented in BBTools (v 39.01)(Bushnell, 2014). Trimmomatic (v 0.39)(Bolger et al., 2014) was used to remove low quality reads and trim reads across samples, which was then followed up with a final read quality assessment using FastQC and MultiQC.

RNA-seq reads were mapped to the *Ca. E. dacicola* genome following indexing of the reference using BWA-MEM and resulting bam files were coordinate sorted and indexed with SAMtools. A gene count matrixed was produced by supplying the indexed .bam files and resulting .gff from PGAP using featureCounts (v 2.0.4)(Liao et al., 2014). FeatureCounts was used with the --countReadPairs, -p, -O, and -M flags to count paired-end reads (specifically count fragments), assign all reads to their overlapping meta-features, and count all multi-mapping reads.

### Pathway Over Representation Analysis (ORA) to identify function affected by pseudogenes

While ORA is traditionally performed on differential abundance data, in this case we directly compared the discrete categories of genes and pseudogenes. ORA was performed using clusterProfiler (v 4.10.1) (Wu et al., 2021) and was done specifically using associated KEGG KO pathways. Statistical testing of association between transposable elements inserted across the *Ca. E. dacicola* genome and pseudogenes

To test if transposable elements (TEs) are more likely to insert into pseudogenes over genes sequences, TEs were first annotated using BLAST+ (Camacho et al., 2009) to the TnCentral database (Ross et al., 2021). Following TE annotation, BEDOPS (v 2.4.41) (Neph et al., 2012) was used to produce input .bed files from the .gff, and the bedtools intersect function of BEDTools (v 2.30.0) (Quinlan & Hall, 2010) was used to determine the TEs that overlapped with genes or pseudogenes coordinates separately. After filtration of all genes and pseudogenes that directly encoded for transposases, a Fisher’s exact test was used to determine if there was nonrandom association between pseudogenes and the presence of TEs.

### Pangenome positive selection analysis

A pangenome positive selection analysis was performed on genomes from *Ca. E. dacicola*, *Ca. E. haradaeae* (aphid-associated symbiont), *E. amylovora* and *E. persicina* (free-living bacterial plant pathogens), and *E. aphidicola* (free-living bacterial plant pathogen that sometimes associated with aphids and thrips) (Table S2). These were a subset of the same genomes downloaded from NCBI and used to conduct phylogenomic analysis. In total, five genomes from each species were used, prioritizing genomes with structural annotations. Analyses were split to either compare *Ca. E. dacicola* to *Ca. E. haradaeae* or *Ca. E. dacicola* to *E. amylovora*, *E. aphidicola,* and *E. persicina*.

Paired protein and transcript coding sequences were extracted from each genome using GffRead (v0.12.7) (Pertea & Pertea, 2020) with each bacterium’s genome FASTA and GFF file. Collectively, these extracted proteins and transcript coding genes were used to create a multi-FASTA index file via CDBtools (v 0.99) (Pertea et al., 2003). For each comparison, the extracted protein sequences were used to predict single copy gene orthologs using OrthoFinder (v 2.5.5). Results from the ortholog prediction were used to create indices using CDBtools and then used to pull out the exact protein and transcript sequences corresponding to each predicted single copy gene ortholog. Protein sequences for each single copy ortholog were aligned with Clustal-Omega (v 1.2.4) (Sievers & Higgins, 2021). With the extracted transcript sequences, codons were mapped back to protein alignments using PAL2NAL (v 14.1)(Suyama et al., 2006). Gene trees were generated for each single copy ortholog codon alignment using RAxML. Branch lengths were removed from each gene tree and only the tree topologies were used. Trees were also modified with a foreground branch designation; for every comparison the foreground branch was the *Ca. E. dacicola* genome.

Positive selection was tested following the methods described in Álvarez-Carretero, as implemented with PAML (v 4.10.7-2)(Álvarez-Carretero et al., 2023). Here, a branch model analysis was employed to identify signatures of positive selection acting on a specific lineage of the phylogenetic tree. Tests were done to evaluate if the selection pressure acting on *Ca. E. dacicola* genes (i.e., the foreground branch) statistically differed from *Ca. E. haradaeae, E. amylovora*, *E. aphidicola,* and *E. persicina* genes (i.e., the background branches), respectively. In the branch model, Ω was allowed to vary on the foreground branch, while Ω was fixed for the background branches. Statistics were calculated by comparing the branch model to the M0 model which estimates one consistent Ω value for the entire gene tree. Extreme ω estimates can arise from poor estimation of dN or dS, particularly when synonymous substitution rates are very low. Therefore, ω values that were below 0.1 or above 3.0 were excluded, as these represented the tails of the global distribution and were likely to reflect estimation artifacts rather than biologically meaningful variation.

## Author contributions

TEH, CJM, and SBM conceived and designed the study, with SMG providing additional advice. CA collected physical specimens, while RLC and TJS performed corresponding wet lab work and created sequencing libraries. TEH conducted the analyses and wrote the manuscript, with CJM, SBM, and SMG providing revision and edits.

## Data availability

The RNAseq data have been deposited with links to BioProject Accession no. PRJNA1291273 in the NCBI BioProject database. Raw reads can be found in the NCBI Sequence Reads Archive: accession numbers SRR34532192, SRR34532191, SRR34532190, SRR34532189, SRR34532188, SRR34532187, SRR34532186, and SRR34532185. Other data and R analysis has been deposited at the USDA NAL Ag Data Commons at https://figshare.com/s/c000bdfa917f57390ce7.

## Funding

Funding for this research is provided through the U.S. Department of Agriculture, Agricultural Research Service appropriated project “Advancing Molecular Pest Management, Diagnostics, and Eradication of Fruit Flies and Invasive Species” (2040-22430-028-000-D).

## Conflict of interest

The authors declare no conflict of interest.

## Acknowledgements

This research employed resources provided by the SCINet project and/or the AI Center of Excellence of the USDA Agricultural Research Service, ARS project numbers 0201-88888-003-000D and 0201-88888-002-000D). The findings and conclusions in this publication are those of the authors and should not be construed to represent any official USDA or U.S. Government determination or policy. Mention of trade names or commercial products in this publication is solely for the purpose of providing specific information and does not imply recommendation or endorsement by the U.S. Department of Agriculture. USDA is an equal opportunity provider and employer. This research was supported in part by an appointment to the Agricultural Research Service (ARS) Research Participation Program administered by the Oak Ridge Institute for Science and Education (ORISE) through an interagency agreement between the U.S. Department of Energy (DOE) and the U.S. Department of Agriculture (USDA). ORISE is managed by ORAU under DOE contract number DE-SC0014664. All opinions expressed in this paper are the author’s and do not necessarily reflect the policies and views of USDA, DOE, or ORAU/ORISE.

## Supporting information

Table S1: Statistics of transcriptomic pair read amounts (in millions) and sizes (Mb) for the olive fruit fly host and *Candidatus Erwinia dacicola* symbiont. Percent read recovery, for symbiont reads, also included.

Table S2: List of *Erwinia* species used to construct a phylogenetic tree. Genbank assembly identification is included for each species. There is also indication of which species were used in the positive selection analysis and effector orthologues analysis.

Table S3: Punatitive predicted effector annotations from DeepSecE including predicted secretion systems and the amino acid length of the related effector sequences.

Table S4: Transposon annotations from TnCentral, including pseudogene status.

Table S5: Table of essential proteins needed for bacterial recombination. There is indication of the presence or absence of these proteins in the *Candidatus Erwinia dacicola* genome, the gene ID they are connected to, and if they are pseudogenes.

Table S6: ORA analysis of significantly enriched pathways between functional genes and pseudogenes.

Table S7: Percent completenesss of associated KEGG pathways in the Candidatus Erwinia dacicola genome.

Table S8: Positive selection analysis comparing single copy orthologues between *Erwinia amylovora, Erwinia aphidicola*, and *Erwinia persicina* each to *Candidatus Erwinia dacicola*. Background dN/dS represents *Erwinia amylovora, Erwinia aphidicola*, and *Erwinia persicina*, while foreground dN/dS represents *Candidatus Erwinia dacicola*.

Table S9: Positive selection analysis comparing single copy orthologues between *Candidatus Erwinia haradaeae* and *Candidatus Erwinia dacicola*. Background dN/dS represents *Candidatus Erwinia haradaeae* while foreground dN/dS represents *Candidatus Erwinia dacicola*.

## Notes

### Competing Interest Statement

The authors have declared no competing interest.

### Summary of Updates

This manuscript has been revised with updated main text, analyses, figures, and supplemental material

